# Timing antigenic escape in multiple myeloma treated with T-cell redirecting immunotherapies

**DOI:** 10.1101/2024.05.22.595383

**Authors:** Marios Papadimitriou, Sungwoo Ahn, Benjamin Diamond, Holly Lee, John McIntyre, Marietta Truger, Michael Durante, Bachisio Ziccheddu, Ola Landgren, Leo Rasche, Nizar J. Bahlis, Paola Neri, Francesco Maura

## Abstract

Recent data highlight genomic events driving antigen escape as a recurring cause of chimeric antigen receptor T-cell (CAR-T) and bispecific T-cell engager (TCE) resistance in multiple myeloma (MM). Yet, it remains unclear if these events, leading to clonal dominance at progression, result from acquisition under treatment selection or selection of pre-existing undetectable clones. This differentiation gains importance as these immunotherapies progress to earlier lines of treatment, prompting the need for innovative diagnostic testing to detect these events early on. By reconstructing phylogenetic trees and exploring chemotherapy mutational signatures as temporal barcodes in 11 relapsed refractory MM patients with available whole genome sequencing data before and after CART/TCE treatment, we demonstrated that somatic antigen escape mechanisms for BCMA- and GPRC5D-targeting therapies are acquired post-diagnosis, likely during CART/TCE treatment. Longitudinal tracking of these mutations using digital PCR in 4 patients consistently showed that genomic events promoting antigen escape were not detectable during the initial months of therapy but began to emerge nearly 1 year post therapy initiation. This finding reduces the necessity for a diagnostic panel to identify these events before CART/TCE. Instead, it underscores the importance of surveillance and identifying patients at higher risk of acquiring these events.

**KEY POINTS:**

1. Genomic events driving antigen escape are recurrent mechanisms of resistance to CART and T-cell engagers in multiple myeloma.
2. Using chemotherapy mutational signatures, we demonstrated that these events are most likely acquired during treatment.

## INTRODUCTION

T-Cell redirecting therapies including chimeric antigen receptor T-cells (CAR-T) and bispecific T-cell Engagers (TCE), have revolutionized the treatment landscape for patients with relapsed/refractory multiple myeloma (RRMM)^1-6^. However, despite these responses, the majority of patients experience relapse within two years of therapy. For the current antigenic targets, B-cell maturation antigen (BCMA) and G protein-coupled receptor class C group 5 member D (GPRC5D), recent studies have shown that genomic alterations of *TNFRSF17* (BCMA) and *GPRC5D* can curtail treatment responses and promote disease progression^7-10^. As recent analyses demonstrate that monoallelic inactivation may be present at baseline and may serve as a risk factor for development of biallelic loss, baseline genomic screening has been proposed to guide the appropriate choice of therapy^11^. However, intrinsic mechanisms of antigen escape may either be selected (i.e., expansion of a pre-existing lesion as a result of a therapeutic bottleneck) or acquired as a result of treatment pressure. While detection of subclonal contributors to resistance before commencement of T-cell redirecting therapy could have significant implications for selecting the most appropriate therapy for RRMM patients, baseline diagnostic testing may prove ineffective in the case of lesions that will be acquired after initiation of therapy. Further hindering our ability to differentiate between these two resistance models (selection or acquisition) are the spatial heterogeneity of MM and the limitations of current sequencing techniques in identifying small subclones^12,13^.

## METHODS

To circumvent these issues, we here developed a novel temporal workflow to time the acquisition of antigenic escape with relation to commencement of T-cell redirecting therapy. We established our workflow using a recently published dataset of 11 RRMM patients with pre- and post-CART/TCE whole-genome sequencing (WGS) data (**Supplementary Methods**)^8^.

## RESULTS AND DISCUSSION

Among these patients, 7 had received anti-BCMA therapy, 1 had undergone anti-GPRC5D therapy, and 3 had received both modalities. Between all patients, a total of 24 WGS samples were analyzed. Of the 11 patients, 7 showed genomic alterations within *TNFRSF17* or *GPRC5D* that were not detectable at baseline using bulk WGS. The key clinical and treatment data for this series are summarized in **Supplementary Table 1**. Our temporal workflow is based on the integrations of two main analyses. In the first analysis DPClust^14^ was used to reconstruct the phylogenetic tree for each patient, distinguishing mutations into truncal (i.e., clonal and shared in all samples), 1^st^-level branch (sample-level events branching from the trunk), and 2^nd^-level branch (sample-level unique branching from the 1^st^-level branch). As expected, all identified mechanisms of antigen escape were localized in the 1^st-^ or 2^nd^-level branches. In the second analysis, since most patients received melphalan-based treatment (n=10/11, e.g., high-dose melphalan and autologous stem cell transplant, HDM-ASCT), we leveraged chemotherapy-associated mutational signatures [e.g., single base substitution (SBS)-MM1/SBS99] as temporal barcodes linked to discrete clinical exposures in each patient’s history (i.e., HDM-ASCT)^15-18^. Specifically, as presence of SBS-MM1/SBS99 in bulk WGS is predicated on the clonal expansion of a single exposed cell bearing a unique melphalan mutational barcode (i.e., single cell expansion model), detection of SBS-MM1/SBS99 only in the phylogenetic trunk implies that *GPRC5D* or *TNFRSF17* somatic escape present in a 1^st^-level branch must have been acquired following the survival and expansion of a single melphalan-exposed cell and therefore was not present before HDM-ASCT (**Supplementary Figure 1**; top). However, the identification of chemotherapy signatures exclusively within the branches in reference to genomic antigen escape events (**Supplementary Figure 1**; middle and bottom) or the absence of any chemotherapy signatures presents a challenge in resolving the temporal estimation. Applying this approach to our series we were able to demonstrate that in 4 patients the mechanisms of antigen escape were not present before HDM-ASCT (i.e. at the time of initial disease diagnosis). Importantly, in two patients we observed evidence of convergent evolution, where disease progressed with at least two clones independently acquiring genomic events on either *TNFRSF17* or *GPRC5D*. In both patients these subclones were localized to the 2^nd^ level branches arising from a 1^st^ level branch undetectable at baseline. Based on the presence of convergent evolution in the 2^nd^ level branches and SBS-MM1/SBS99 melphalan signatures in the trunk, it is likely that these intrinsic mechanisms of antigen escape were acquired and subsequently expanded under CART/TCE selective pressure. Below we summarize these cases in more details (**Figure 1** and **Supplementary Figure 2**).

**Figure 1:**
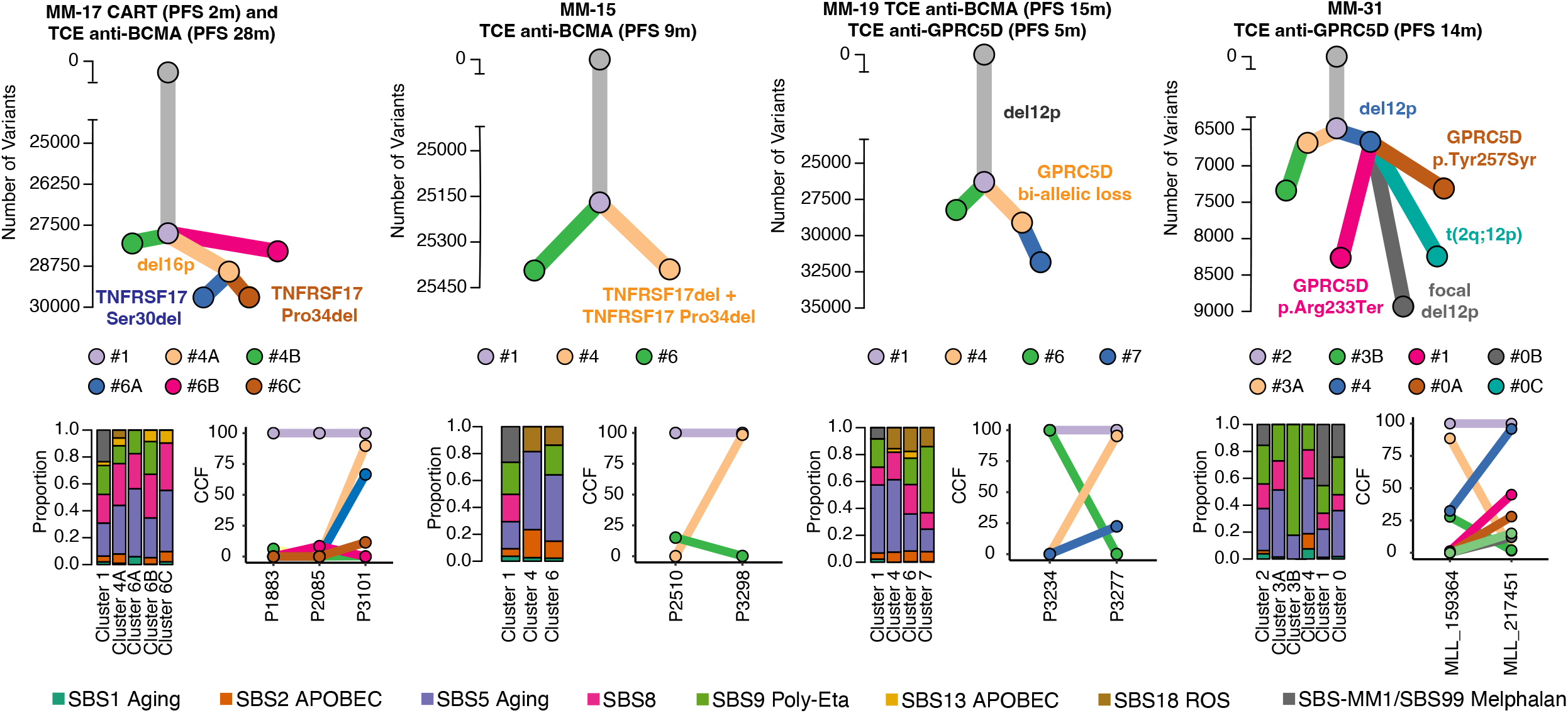
Timing antigen escape mechanisms in RRMM treated with CART/TCE. Selected cases for timing the acquisition of TCE/CART antigen escape mutations. Clusters in the phylogenetic tree were reconstructed using DPClust. Branch length is proportional to the number of single base substitution (SBS) in each cluster. Antigen escape variants are color-coded according to the cluster in which they are found. Mutational signatures analyses were run on clusters with >50 SBS. For example, Cluster #4B in MM-17 had 7 mutations and so mutational signature fitting was not performed. In MM-31 Cluster #0 grouped all the subclonal variants detected at relapse after TCE anti-GPRC5D. Presence of convergent evolution with 4 different and independent subclones was previously demonstrated by phasing reads.^8^ This is why the cluster appeared split in the three but is reported as one for the SBS analysis. PFS, Progression-Free Survival.

Patient MM-17 was treated with both anti-BCMA CART and TCE. Notably, short progression-free survival (PFS) following CART was likely due to product qualitative manufacturing failure. After this the patient achieved a prolonged response following anti-BCMA TCE and progressed after 24 months. Three samples were investigated by WGS: P1883 (pre-CART), P2085 (post-CART), and P3101 post-TCE. Six major clusters were identified: #1, #4A, #4B, #6A, #6B, and #6C. Cluster #1 included all clonal SBS shared between the samples (i.e. trunk). Clusters #4B and #6B were detected at subclonal level only in the samples collected before and after CART, respectively. Cluster #4A, which harbors a monoallelic deletion 16p, was undetectable in the pre-TCE and pre-CART samples, however it became clonal post-TCE. From this cluster, two 2^nd^ level branches emerged: Cluster #6A and Cluster #6C carrying two independent mutations in the *TNFRSF17* extracellular domain (CCF of 90% and 10%, respectively, i.e. convergent evolution). Based on our temporal model, because SBS-MM1/SBS99 was only detected in the trunk, concurrent with convergent evolution, we can conclude that two were acquired, and not present before HDM-ASCT (or TCE). Importantly, the absence of these events before treatment with CART/TCE was also validated in digital PCR for the two *TNFRSF17* mutations.

Patient MM-15 had samples collected before and after anti-BCMA TCE. Cluster #1 was clonal in both samples (i.e., trunk of phylogenetic tree). Cluster #4 was subclonal before TCE and not detectable at progression, while Cluster #6 was not detectable before treatment but clonal at progression. This subclone carries monoallelic focal loss of *TNFRSF17* coupled with a clonal *TNCFRSF17* Pro34del. Presence of SBS-MM1/SBS99 in the trunk reveals that both BCMA somatic events were not present at the time of HDM-ASCT.

Patient MM-19 underwent four lines of therapy. Notably, they received melphalan exposure during ASCT and subsequently received both anti-GPRC5D (talquetamab) and anti-BCMA TCE (elranatamab) treatments. This patient also had single-cell copy number variant-sequencing (scCNV-seq) on bone marrow plasma cells before and after talquetamab therapy. Additionally, bulk WGS was performed on two distinct samples both after relapse on talquetamab treatment (P3234 and P3277). We identified four distinct phylogenetic clusters among the relapsed samples (#1, #4, #6, #7). Cluster #1 encompassed all clonal SBS present in both samples, indicative of a common ancestry. Interestingly, the patient exhibited evidence of branching evolution, with Cluster #4 and #6 emanating from Cluster #1. In the second relapse sample (P3277), Cluster #4 attained clonality, while cluster #6 diminished. In this sample, Cluster #4 (CCF=0.95) contained a biallelic *GPRC5D* loss and presence of SBS-MM1/SBS99 within the truncal SBS suggests that loss of antigen occurred following melphalan exposure.

Patient MM-31 received six lines of treatment and was twice exposed to melphalan (tandem HDM-ASCT), followed by bendamustine (which causes a similar mutational signature profile to melphalan^19^). They later received anti-GPRC5D TCE treatment with bulk WGS samples taken pre- and post-therapy. Subclonal reconstruction identified 6 major clusters (#0, #1, #2, #3A, #3B, #4). Cluster #2 contained all clonal SBSs with branching evolution leading to Clusters #3a, which diminished after treatment, and #4, which became clonal. The relapsed sample showed a large deletion in the short arm of chromosome 12 (12p) encompassing *GPRC5D* within Cluster #4. Profound convergent evolution then occurred with 4 distinct clusters arising from Cluster #4, each with its own second hit on *GPRC5D* leading to biallelic inactivation. Cluster #1 harbored a nonsense mutation (p.Arg233Ter) with a CCF of 0.45; #0A contained a missense mutation in *GPRC5D* with a CCF of 0.28; #0B harbored a second, focal deletion of *GPRC5D* (CCF=0.12); while #0C contained a reciprocal translocation involving the long arm of chromosome 2 and the short arm of chromosome 12 (CCF=0.15). Here, SBS-MM1/SBS99 was truncal, corresponding to a single-cell expansion after HDM-ASCT. Because SBS-MM1/SBS99 is also present in the 2^nd^ level relapse branches and not the 1^st^-level branches, this later SBS-MM1/SBS99 likely is related to bendamustine exposure^19^ and therefore implicates the 2^nd^-level convergent subclones as having acquired their respective antigen-escape mutations after HDM-ASCT exposure. Consequently, in MM-31, it can be inferred that all *GPRC5D* negative clones were not originally present at the time of initial diagnosis.

To further validate these findings, for 4 out of 11 patients, we conducted digital PCR (ddPCR) at various time points using bone marrow samples collected before, during, and after treatment with TCE (**Figure 2** and **Supplementary Methods**). Two of these patients had SBS-MM1/SBS99 (MM-17 and MM-15, **Figure 1**). All experiments consistently demonstrated that genomic events promoting antigen escape were undetectable by ddPCR prior to and during the initial months of TCE therapy but rather emerged later on up to 4 months prior to measurable biochemical or clinical relapse. Overall, this strongly supports the hypothesis that these events are acquired rather than selected.

**Figure 2:**
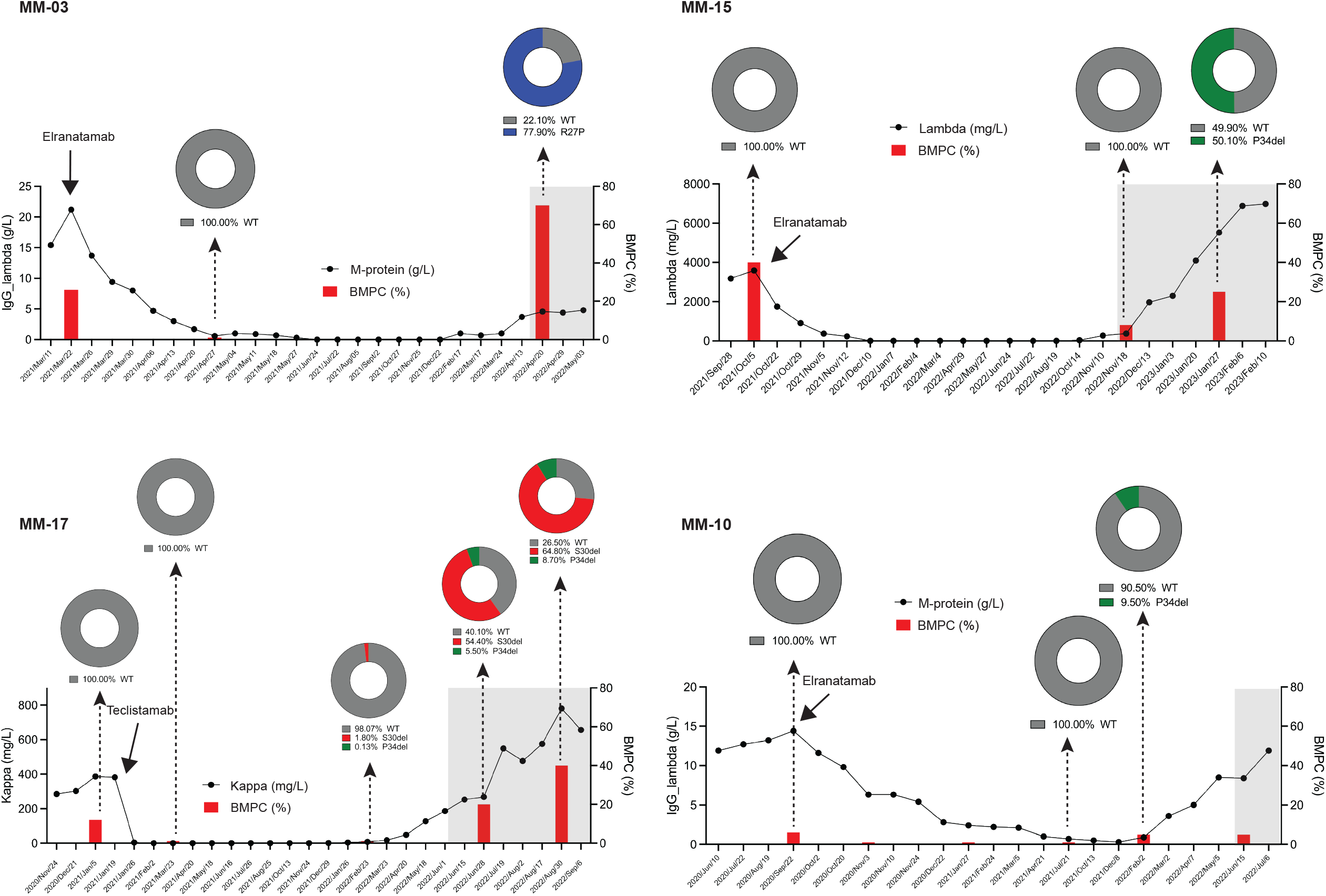
Tracking onset of mutations on *TNFRSF17* over time in RRMM treated with CART/TCE. ddPCR reveals the emergence of *TNFRSF17* mutants at later time points post TCE therapy initiation. Plots depict the serial measurements of the serum M-protein by serum electrophoresis or free light chain studies (solid black line), the percent of the bone marrow plasma cells (BMPC %, red bars), the donut plots indicate *TNFRSF17* WT and mutants VAF and the grey zone reflects the time of biochemical or clinical relapse.

Defining the molecular clock of mutations in antigens targeted by T-cell adaptive therapies has large prognostics and therapeutic implications and may dictate the utility of prescreening versus surveillance for these mutations as well as the benefit and timing of dual versus sequential T-cell targeted approaches. Our temporal workflow, which integrates clonal phylogeny with chemotherapy mutational signatures as temporal barcodes, reveals that somatic mechanisms of antigen escape for T-cell redirecting therapies that target BCMA and GPRC5D were not present at diagnosis for at least 4 of 7 patients. These lesions were acquired, most likely under the selective pressure of CART/TCE. For the remaining patents, our workflow could not establish when these events occurred. These findings hold significant importance, especially concerning GPRC5D,^2,8,10,20^ where antigen escape appears to be the primary mechanism of resistance in both CART and TCE therapies. While their relevance for anti-BCMA therapies is predominantly associated with TCE,^21^ it remains unclear whether the acquisition of the same BCMA mutations identified after TCE treatment play a significant role in post-CART settings. Overall, our observation aligns with the general absence of these genomics events across a large cohort of newly diagnosed MM cases with available whole-exome sequencing (WES) or WGS data (n=752). The demonstration that antigen escape mechanisms may be acquired somatic events – rather than being selected from pre-existing subclones – diminishes the need for developing a diagnostic panel for their detection prior to CART/TCE treatment. Instead, it underscores the need to investigate which patients are at a higher risk of acquiring these events – and whether patients might benefit from an alternate CART/TCE following relapse – especially as CART/TCE enter earlier lines of therapy for MM.

## DATA SHARING STATEMENT

All WGS data will be uploaded on EGA.

## ACKNOWLEDGEMENTS

This work was supported by the Myeloma Solutions Fund (MSF), Paula and Rodger Riney Multiple Myeloma Research Program Fund, the Tow Foundation, Sylvester Comprehensive Cancer Center NCI Core Grant (P30 CA 240139).

FM is supported by the American Society of Hematology (ASH), Leukemia & Lymphoma Society (LLS), and by International Myeloma Society (IMS).

B.D. is supported by the Sylvester K12 Calabresi Clinical Oncology Research Career Development Program.

## AUTHORSHIP CONTRIBUTIONS

N.J.B., L.R., and F.M. conceived, designed and supervised all experiments, performed the analysis, and wrote the paper. H.L., M.P., S.A., and B.D. analyzed the data, performed analyses, and wrote the paper. M.C. performed has performed the digital PCR; M.T., and O.L. contributed to patient consent and sample collection. M.D. and B.Z. analyzed the data and performed analyses.

## SUPPLEMENTARY FIGURE LEGEND

**Supplementary Figure 1: Selection vs. acquisition of antigen escape mechanisms**. Cartoons depicting multiple myeloma tumor evolution with respect to high-dose melphalan-autologous stem-cell transplant (HDM-ASCT), TCE/CART therapy, and bulk WGS. SBS-MM1/SBS99 (dark red) in the trunk of the tumor phylogenetic tree (top) implies that a single-cell expansion took place following HDM-ASCT leaving a clonal melphalan barcode and that subsequent antigen escape mutations were acquired in downstream phylogenetic branches. SBS-MM1/SBS99 fit to the branches of the tree do not provide the resolution to detect acquisition (middle) vs. selection (bottom). TCE, T-Cell Engager; CART, chimeric antigen receptor T-cell; WGS, whole genome sequencing; SBS, single base substitution.

**Supplementary Figure 2:** Cases for which antigen escape timing was not possible. Branch length is proportional to the number of SBS in each cluster. Antigen escape variants are color-coded according to the cluster in which they are found. Mutational signatures analyses was run on clusters with >50 SBS. In MM-02, MM-10 and MM-18 the timing of mechanisms of antigen escape was not possible because SBS-MM1 was present in both trunk and branches of the phylogenetic tree. In MM-03 no SBS-MM1/SBS99 signature was detected. In MM-04, MM-06 and MM-20 no mechanisms of antigen escape was detected. In MM-02, Cluster 3A carried a biallelic loss of *TNFRSF17B* and was detected by single cell CNV before therapy (Lee et al. Nat Med 2023).

