## Supplementary figures and images for "Timing antigenic escape in multiple myeloma treated with T-cell redirecting immunotherapies"

### Supplementary Figure 1

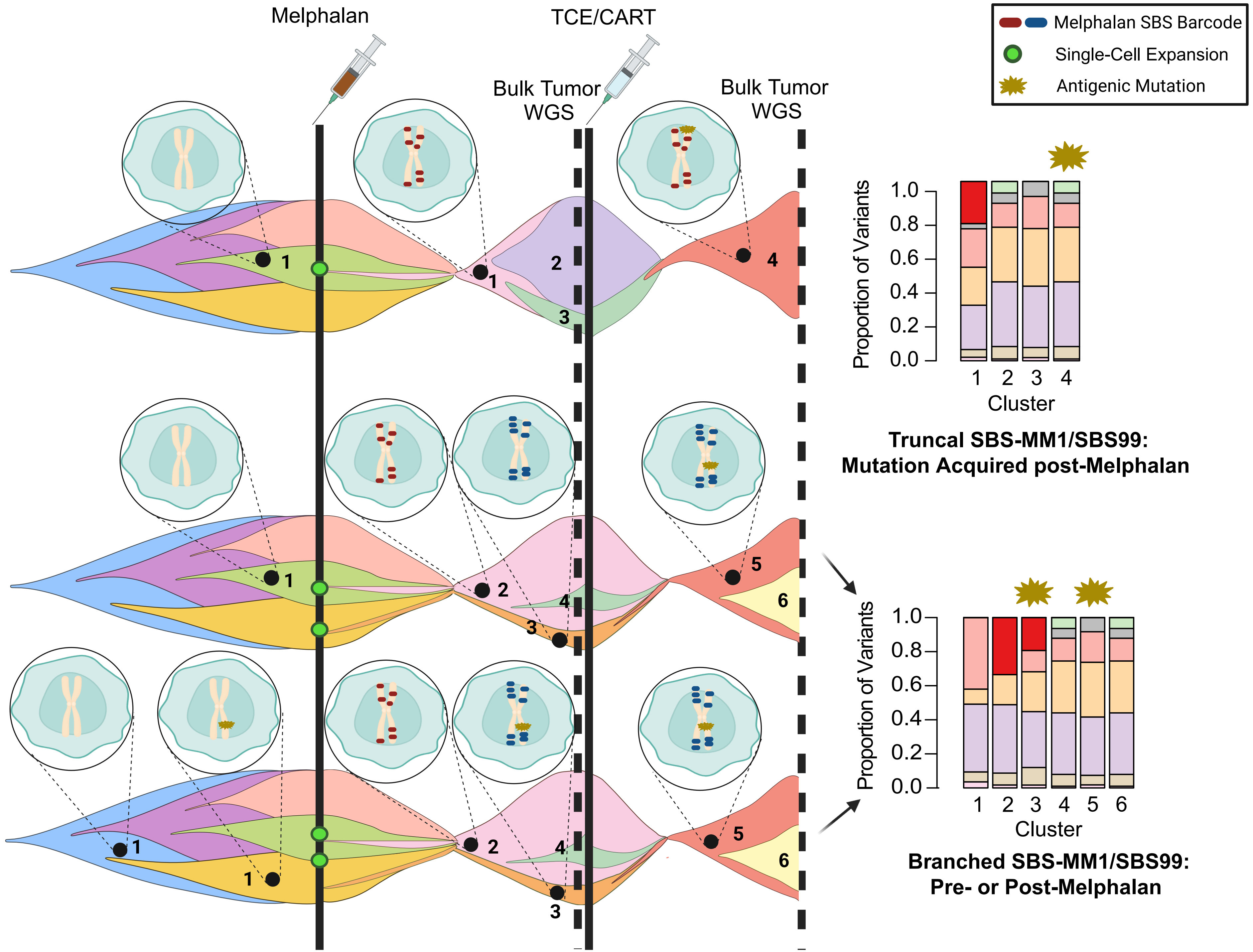

### Supplementary Figure 2

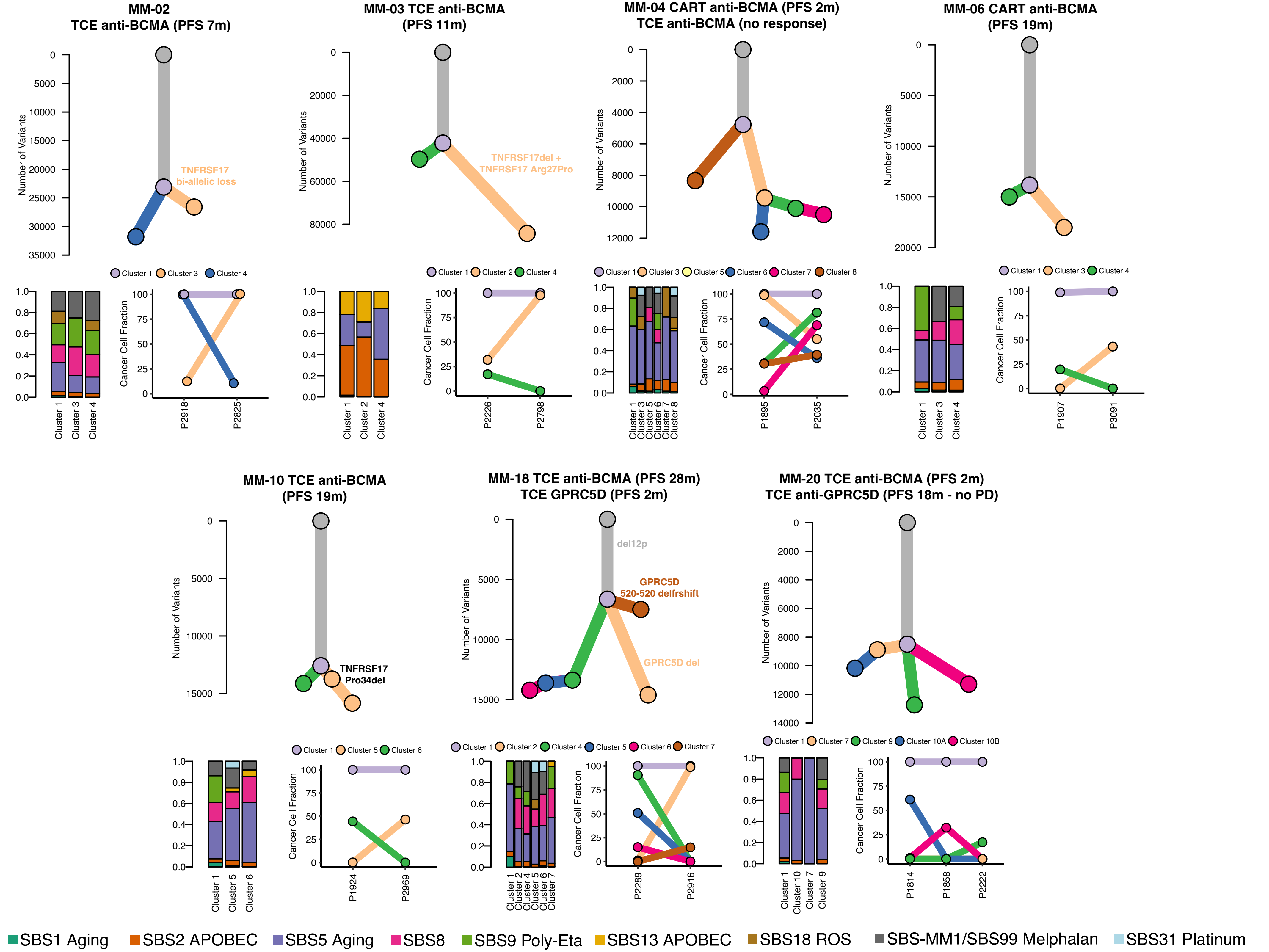
