## Supplementary Methods for "Timing antigenic escape in multiple myeloma treated with T-cell redirecting immunotherapies"

**Patient CD138^+^ sample collection**

This study, adhering to the Declaration of Helsinki, received approval from the University of Calgary's conjoint health research ethics board (HREBA.CC-21-0248) and the University Hospital of Würzburg's ethics board (Würzburg EK 8/21). Patient participation was contingent on written informed consent. The study involved 11 multiple myeloma (MM) patients treated under various protocols, including treatments with anti-BCMA CAR T/TCE and anti-GPRC5D CAR T/TCE, along with other non-anti-BCMA/GPRC5D treatments. The cohort comprised patients treated with anti-BCMA (n=7), anti-GPRC5D (n=1), or both (n=3). CD138^+^ MM cells were isolated from patient bone marrow aspirates for subsequent analysis, including single-cell RNA (scRNA-seq) and copy number changes (scCNV-seq) sequencing, as well as bulk whole genome sequencing (WGS). Study sample set clinical summary is reported in **Supplementary Table 1**.

**WGS**

DNA from CD138+ myeloma cells underwent WGS using protocols tailored to the sample collection site. For samples collected at the University of Calgary, the New York Genome Center (NYGC) performed sequencing, employing the TruSeq DNA Nano Library Preparation Kit, while at the University Hospital Center Würzburg, the TruSeq PCR-free library prep kit was used. Sequencing was done on Illumina Novaseq 6000 sequencer. and the subsequent data analysis utilized a suite of different calling tools, such as *Mutect2*, *Strelka* and *Lancet* for SNVs and indels; *GATK4* and *ASCAT* for copy number changes, as described previously^8^.

**ScRNA-seq and scCNV-seq analysis**

Using the 10x Genomics system, CD138+ myeloma cells underwent scRNA-seq and scCNV-seq, with cells encapsulated in gel beads for precise barcode and primer integration, followed by cDNA and gDNA amplification. Quality control preceded sequencing on an Illumina NextSeq 500. Data were analyzed with the CellRanger suite and the Seurat package in R for gene expression, while custom scripts assessed CNV data, streamlining the interpretation of complex genomic variations. Details on the pipeline implemented for single cell analysis have been described previously^8^.

***TNFRSF17* and *GPRC5D* mutations in newly diagnosed multiple myeloma.**

To define the prevalence of *TNFRSF17* and *GPRC5D* changes, we interrogated 752 newly diagnosed multiple myeloma (NDMM) patients from the CoMMpas cohort. This analysis revealed, at baseline, monoallelic loss of *TNFRSF17* or *GPRC5D* in 21 (2.79%) and 68 (9.04%) samples, respectively. No patient carried any biallelic loss in either of the genes. Furthermore, 7 samples were found to have mutational events on *TNFRSF17*, while only 2 samples carried a mutation on *GPRC5D* at baseline.

**Phylogenetic reconstruction.**

To determine the tumor clonal architecture, we used *DPClust* (<https://github.com/Wedge-lab/dpclust>). As has been described, phylogenetic trees were resolved according to the Pigeonhole principle. To provide enhanced subclonal resolution on a case-by-case basis, as described in the manuscript, further splitting of clusters was performed based on the presence of scCNV-seq and read phasing (to demonstrate mutual exclusivity of variants). The mutational signature analysis of single-base substitutions (SBSs) was performed using *sigprofiler* (<https://github.com/AlexandrovLab/SigProfilerExtractor>) and *mmsig* only in clusters with >50 mutations (<https://github.com/evenrus/mmsig>). All analyses were performed in R (v4.3.1).

**Digital PCR validation**

TNFRSF17 mutations were screened on genomic DNA extracted from sorted CD138 bone marrow plasma cells using the QuantStudio 3D Digital PCR system (Thermo Fisher Scientific). Genotyping assays were designed using the customized TaqMan Assay Design Tool with designed primers and probes sequences as previously published (Lee et al. Nat Med 2023). For dPCR reactions genomic DNA (10 ng) were loaded on to a QuantStudio 3D Chip v.2 on a ProFlex Flat PCR thermocycler. The following thermocycler conditions were used: 96 °C, 10 min (60 °C, 2 min and 98 °C, 30 s) ×39 cycles, 60 °C, 2 min. QuantStudio 3D dPCR Instrument was used for Chip reading and analysis performed using QuantStudio 3D AnalysisSuite Software v.3.1.6.
